# Urinary extracellular vesicles reveal a sex-specific miRNome profile in alcohol use disorder patients

**DOI:** 10.64898/2026.07.08.737166

**Authors:** Blanca Martín-Urdiales, Carla Perpiñá-Clérigues, Susana Mellado, Maura Rojas-Pirela, María-Lourdes Aguilar Sánchez, David Puertas-Miranda, Francisco García-García, Miguel Marcos, María Pascual

**Affiliations:** Department of Physiology, School of Medicine and Dentistry, University of Valencia, 46010 Valencia, Spain; Computational Biomedicine Laboratory, Príncipe Felipe Research Center, 46012 Valencia, Spain; Department of Internal Medicine, University Hospital of Salamanca, 37007, Salamanca, Spain; Institute of Biomedical Research of Salamanca (IBSAL), 37007 Salamanca, Spain; Primary Care Addiction Research Network (RIAPAd), Instituto de Salud Carlos III, 28029, Madrid, Spain; Department of Medicine, University of Salamanca, 37001, Salamanca, Spain; Psychiatry Service, University Hospital of Salamanca, 37007, Salamanca, Spain

**Author notes:** Corresponding author: María Pascual, Department of Physiology, University of Valencia, Avda. Blasco Ibáñez, 15. 46010 Valencia, Spain. These authors contributed equally to this work (Blanca Martín-Urdiales, Carla Perpiñá-Clérigues).

**Keywords:** miRNA transcriptomics, extracellular vesicles, urine, alcohol use disorder, sex-based differences

## Abstract

miRNA-based transcriptomic analysis of extracellular vesicles (EVs) provide a promising strategy for identifying non-invasive biomarkers and understanding complex pathological mechanisms. Recently, however, urinary extracellular vesicles (uEVs) have emerged as a valuable window into molecular alterations. Despite the high morbidity and mortality associated with alcohol use disorder (AUD), the molecular mechanisms underlying its sex-specific differences remain poorly understood. To address this, we characterize for the first time the uEV miRNome in AUD, revealing its sexually dimorphic profile. We employed uEVs from actively drinking AUD patients of both sexes who did not have advanced liver disease, alongside matched controls. Deep sequencing revealed 14 differentially expressed miRNAs in females (e.g., hsa-miR-197-3p, hsa-miR-19b-3p, hsa-miR-505-3p, hsa-miR-625-5p, and hsa-miR-27a-5p) and 6 in males (e.g., hsa-miR-1290, hsa-miR-1246, hsa-miR-450a-5p, and miR-590-5p). Notably, whereas hsa-miR-4787-5p was consistently overexpressed in uEVs from both sexes, it was absent in plasma-derived EVs, highlighting the specificity of the urinary compartment. Remarkably, the miRNA signatures we uncovered reflect the multiorgan impact of AUD. For instance, hsa-miR-1290 and hsa-miR-197-3p point to alcohol-related liver injury and systemic inflammation, whereas hsa-miR-19b-3p and hsa-miR-1246 signal neuroinflammation and neuronal stress. A subset—including hsa-miR-1290, hsa-miR-1246, and hsa-miR-27a-5p—has been implicated in cancer contexts. Collectively, these findings support the uEV miRNome as a promising sex-informed molecular signature of AUD with biomarker and mechanistic relevance.

## Introduction

MicroRNAs (miRNAs) are short noncoding RNAs of approximately 22 nucleotides that regulate gene expression at the post-transcriptional level through mRNA interference mechanisms [1, 2]. Since individual miRNAs can regulate multiple mRNAs, and each mRNA can be regulated by several miRNAs, dysregulation of these molecules can disrupt cellular homeostasis and contribute to the development of diverse pathological conditions [3, 4]. Recent advances in transcriptomic approaches, especially high-throughput RNA sequencing (RNA-seq), have facilitated comprehensive profiling of miRNA expression, offering robust tools for biomarker identification and for unraveling the molecular mechanisms underlying diverse pathologies [5, 6]. Thus, miRNA-based transcriptomic analysis represents a robust and sensitive approach to uncover molecular alterations associated with complex disorders.

Extracellular vesicles (EVs) are nanosized particles that carry molecular cargo, including miRNAs, reflecting the physiological and pathological state of their cells of origin [7, 8]. Analyzing their content offers a valuable, non-invasive approach to investigate molecular alterations associated with various diseases, including neurodegenerative disorders [9, 10]. Among EVs, urinary EVs (uEVs) have attracted particular interest as a non-invasive source of biomarkers, since their molecular cargo can reflect biological processes both within the urinary tract and in systemic conditions, including neurodegenerative diseases and cancer [11, 12]. Therefore, the analysis of miRNA content in uEVs represents a promising approach to identify miRNA biomarkers useful for diagnosis, monitoring, and mechanistic studies across diverse pathological conditions [13, 14].

Alcohol use disorders (AUD) encompass a heterogeneous spectrum of clinical manifestations, resulting from excessive alcohol consumption [15] and account for nearly three million deaths worldwide each year, representing a major cause of morbidity and mortality [16]. AUD is associated with diverse health consequences, including alcohol-related liver disease, hepatocellular carcinoma, cardiovascular complications, and neuropsychiatric disorders, with the brain and liver being the most affected organs [17]. Recent transcriptomic studies have highlighted a growing interest in identifying molecular biomarkers—particularly non-invasive ones—to assess prior alcohol exposure and the severity of alcohol-induced organ damage [18, 19]. In this context, circulating EVs and their molecular cargo have recently emerged as promising biomarkers for detecting ethanol exposure and its systemic effects [20, 21]. Therefore, this study pioneers the analysis of uEV miRNA profiles in female and male AUD patients to identify signatures associated with sex and alcohol-related pathophysiology. We identified distinct uEV miRNA profiles in females (14 miRNAs) and males (6 miRNAs) with AUD, including the urine-specific hsa-miR-4787-5p. These miRNAs map onto key AUD pathologies—liver injury, neuroinflammation, and cancer-related processes—establishing the uEV miRNome as a sex-informed molecular signature with biomarker potential.

## Material and methods

### Human subjects and experimental design

A total of 11 patients (7 males and 4 females) with AUD, diagnosed according to DSM-5 criteria, were recruited for this study through the Alcoholism Unit of the University Hospital of Salamanca (Spain). The mean age of the male AUD group was 48 years, and that of the female AUD group was 47 years. All patients reported active alcohol consumption of ≥100 g ethanol/day prior to study enrollment. Although it was difficult to determine the exact timing of the last alcohol intake, none of the patients had consumed alcohol immediately before sample collection, thereby ruling out a significant contribution from acute ethanol ingestion. Laboratory tests showed normal values for prothrombin time, hemoglobin concentration, and serum albumin levels. All patients tested negative for hepatitis B surface antigen and antibodies against hepatitis C virus. None of the participants presented acute or chronic medical conditions that could interfere with the study outcomes, nor were they polydrug users. Advanced liver disease was excluded by clinical examination, laboratory analyses, and ultrasound assessment. Patients presenting physical signs of chronic liver disease (e.g., cutaneous stigmata, hepatomegaly or splenomegaly, gynecomastia, testicular atrophy, and/or muscle wasting), ultrasonographic abnormalities beyond hepatic steatosis, or serum transaminase levels exceeding 2–3 times the reference range were excluded (Table S1). A group of 7 healthy male volunteers and 6 healthy female volunteers who reported alcohol consumption of <15 g ethanol/day were included as controls, with mean ages of 47 and 52 years, respectively. All participants provided written informed consent, and the study protocol was approved by the Ethics Committee of the University Hospital of Salamanca, Spain (Project ID: PI 2023 07 1389).

Urine samples were collected in sterile, additive-free 50 mL polypropylene tubes and stored at −80 °C until further processing for EV isolation. The study design, aimed at examining alcohol-induced changes in uEV miRNA content and potential sex-specific molecular responses, is summarized in Figure 1. miRNA profiling from uEVs included four major steps: (i) extraction and sequencing of EV-derived miRNAs; (ii) preprocessing, mapping, and normalization; (iii) differential expression analysis; and (iv) functional enrichment analysis.

**Figure 1.**
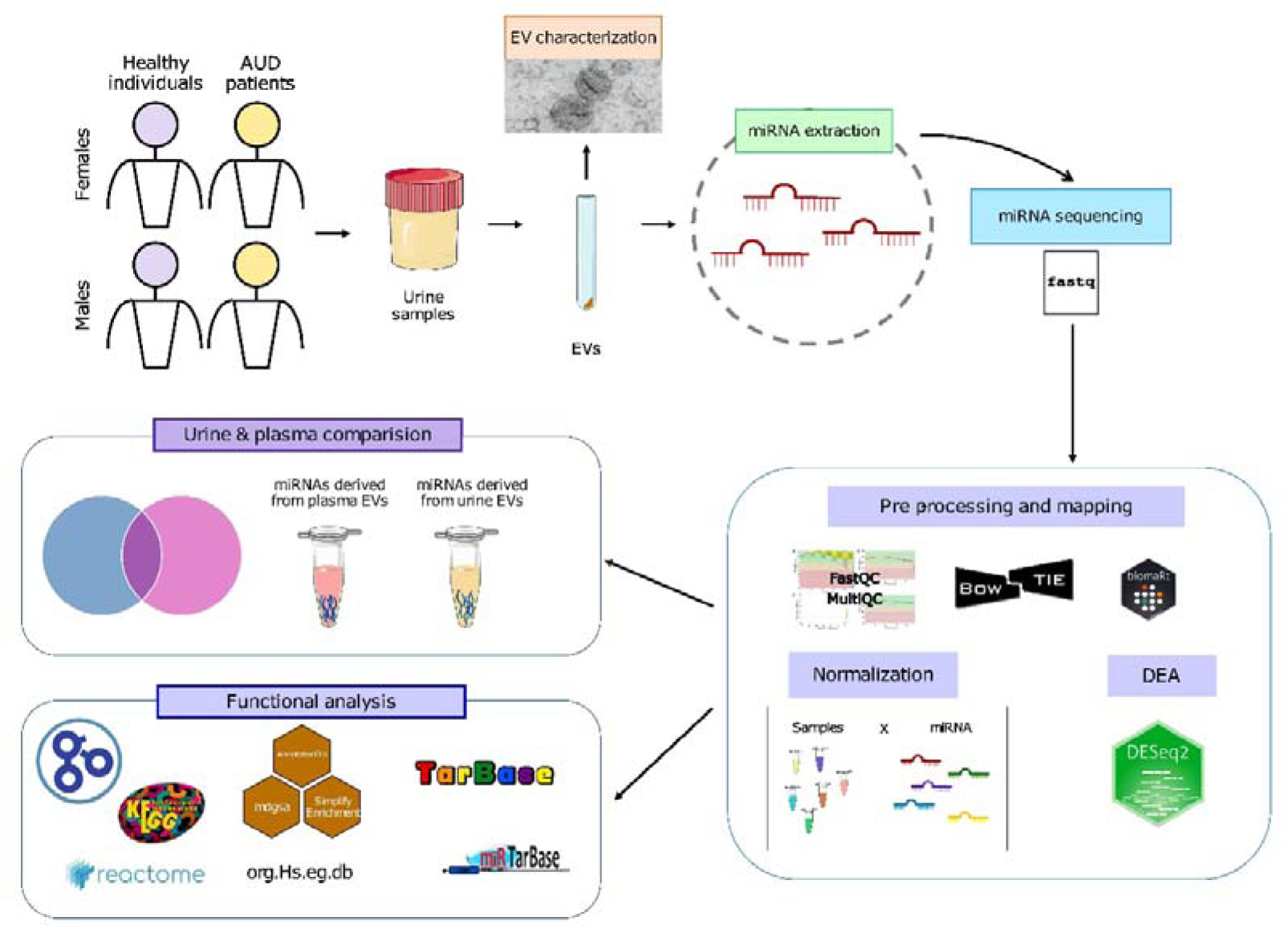
Experimental design and workflow. miRNAs were extracted from uEVs obtained from healthy individuals and alcohol use disorder (AUD) patients of both sexes. After data normalization, differential expression analysis (DEA) was performed on the miRNA profiles, along with the identification of miRNAs shared between the uEVs and our previously published plasma EV data from AUD patients [22]. Functional analyses were subsequently conducted based on these results.

### EV isolation from human urine

Urine samples were first centrifuged at 300 × g for 10 min to remove residual cells and debris. The resulting supernatant was subjected to a second centrifugation at 2,000 × g for 10 min, transferred to new tubes, and subsequently ultracentrifuged at 100,000 × g for 2 h at 4 °C. The pellet containing EVs was washed with phosphate-buffered saline (PBS) and centrifuged again at 100,000 × g for 1 h to obtain purified EVs. The final pellet was used for subsequent EV characterization and miRNA extraction.

### uEV characterization by transmission electron microscopy, western blot, and nanoparticle tracking analysis

uEVs were resuspended in PBS and characterized by transmission electron microscopy (TEM), Western blotting, and nanoparticle tracking analysis (NTA), as previously described [23]. For TEM, EVs morphology were examined using an FEI Tecnai G2 Spirit microscope (FEI Europe, Eindhoven, The Netherlands) with a Morada digital camera (Olympus Soft Image Solutions GmbH, Münster, Germany). Western blot analyses were performed using antibodies against CD9, CD63, CD81, and calnexin (Santa Cruz Biotechnology, USA). Figure S1 shows representative blots for each marker. The absolute size range and concentration of EVs were determined by NTA using a NanoSight NS300 instrument (Malvern Panalytical, Malvern, UK).

### RNA extraction and sequencing

Total RNA was extracted from uEVs using the Total Exosome RNA Isolation Kit (Invitrogen, Waltham, MA, USA) according to the manufacturer’s protocol. miRNA libraries were prepared from total RNA using the NEXTFLEX® Small RNA-Seq Kit v4 with UDIs (Revvity Inc., Massachusetts, USA). Following the manufacturer’s protocol, briefly: small RNA molecules were first ligated to 3’-4N adenylated adapters, taking advantage of the phosphate group at their terminal end, which allows exclusive selection of these molecules. Subsequently, 5’-4N adapters were ligated. After adapter ligation, the molecules were reverse-transcribed into cDNA. The resulting cDNA fragments were amplified and indexed by PCR using primers with unique barcodes for each sample. Finally, a size-selection purification was performed. Library quality control and concentration were assessed using the Agilent Technologies 2100 Bioanalyzer with high-sensitivity DNA chips. An equimolar pool of each library was then prepared for sequencing on the NextSeq 550 NGS platform (Illumina, San Diego, CA, USA) using 50-cycle single-read sequencing (1 × 50 bp). The raw sequencing output was subsequently exported in fastq format for downstream analysis.

### Processing and mapping

Raw sequencing reads were processed with Cutadapt v4.6 [24] to remove the Illumina adapter sequences and filtered based on length (17–35 nt) and quality (Phred score ≥ 30). The resulting reads were aligned against a curated reference database of mature *Homo sapiens* miRNA sequences from miRBase (mature.fa) [25] using Bowtie2 v2.5.3 [26]. Alignment parameters (-L 6 -i S,0,0.5 --ignore-quals -norc --score-min L,-1,-0.6 -D 20) were adapted from Locati et al. [27], optimizing sensitivity and specificity for small RNA alignment. Read counts were obtained directly from the SAM output files produced during alignment. Distinction between uniquely and multi-mapped reads was performed using Bash grep commands, and only those reads that mapped uniquely to the reference were kept for downstream quantification. This procedure reduces potential biases caused by ambiguous mappings and ensures reliable assessment of miRNA expression levels.

### Exploratory analysis and normalization of miRNA profiles

Data filtering, normalization, and all bioinformatic and statistical analyses were performed in R (version 4.3.2) [28] using custom scripts. miRNAs were retained if they showed at least 1 count per million (CPM) in four or more samples. Normalization was performed using the median ratio method implemented in DESeq2 [29] to correct for variations in sequencing depth and RNA composition among samples. Exploratory analyses—including boxplots, principal component analysis (PCA), and hierarchical clustering—were conducted both prior to and following normalization to assess sample distribution across experimental groups and detect potential outliers or batch effects.

### Differential expression analysis of miRNAs

Differentially expressed miRNAs were identified using the DESeq2 package [29]. P-values were adjusted for multiple testing using the Benjamini–Hochberg (BH) method [30], and miRNAs with an adjusted *p*-value ≤ 0.05 were considered statistically significant. Three primary comparisons were performed to assess sex-specific effects in AUD: (i) Impact of AUD in females (**IF**): comparison between female AUD patients and female controls (*AUD.Females − Control.Females*), (ii) Impact of AUD in males (**IM**): comparison between male AUD patients and male controls (*AUD.Males − Control.Males*), and (iii) Impact of sex in AUD (**IS**): comparison between IF and IM [(*AUD.Females − Control.Females) − (AUD.Males − Control.Males*)].

The IF and IM analyses identified miRNAs altered by AUD within each sex, whereas the IS comparison revealed miRNAs whose responses to AUD differed significantly between males and females, accounting for baseline differences observed in healthy individuals. Expression changes were quantified as log fold change (LFC), where the absolute value reflects the magnitude of the change and the sign indicates directionality. A positive LFC denotes higher expression in the first group (AUD groups in IF/IM), while a negative LFC indicates higher expression in the second group (controls in IF/IM). For the IS contrast, LFC reflects the interaction between sex and AUD, indicating whether the AUD-related change in expression is more positive (or more negative) in females compared to males.

### Functional profiling of miRNAs

To investigate the functional implications of miRNA dysregulation, differentially expressed miRNAs were mapped to their target genes using two curated databases: TarBase [31] and miRTarBase [32]. Subsequently, the mdgsa package (v.0.99.2) [33] was used to transfer differential expression statistics from miRNAs to their corresponding target genes. This procedure considers both statistical significance (adjusted p-values) and the direction of change (LFC sign), generating a gene-level ranking that reflects the average regulatory effect of multiple miRNAs on a single target gene.

Functional characterization was performed using a gene set enrichment analysis (GSEA) approach to identify biological processes associated with miRNA-driven gene expression changes. Gene Ontology (GO) terms [34] —including Biological Process (BP), Cellular Component (CC), and Molecular Function (MF)—as well as Reactome [35] and Kyoto Encyclopedia of Genes and Genomes (KEGG) [36] pathways were evaluated. GO annotations were obtained using org.Hs.eg.db (v.3.18.0) [37] and GO term definitions were retrieved from the GO.db package [38]. Reactome annotations were obtained using reactome.db [39], while KEGG pathway identifiers were assigned using org.Hs.eg.db and annotated using the KEGGREST package [40].

P-values were adjusted for multiple testing using the Benjamini–Hochberg (BH) method [30], and GO terms or pathways with an adjusted p-value ≤ 0.05 were considered significant, indicating biological processes or pathways potentially affected by miRNA-mediated regulation.

Significant terms and pathways (BH-adjusted p ≤ 0.05) were prioritized for interpretation. Visualization of GO terms was performed using the binary_cut function from simplifyEnrichment [41] to group terms based on semantic similarity. Additionally, dot plots were generated for specific terms and pathways to enhance the interpretation of the miRNA regulatory network in the context of disease pathology.

### Web platform

An interactive web platform (https://bioinfouvcipf.shinyapps.io/miRNAuEVs-AUD/) was developed to provide comprehensive documentation of the computational methodologies used in this study, built with the R Shiny framework (v.1.12.1) [42]. This resource allows users to explore the complete dataset and interactively access differential miRNA expression results and functional analyses of interest. It also provides access to additional analyses and visualizations not included in the main manuscript. As an open-access tool, the platform facilitates data sharing, supports innovative research, and promotes the discovery of novel findings in the field.

## Results

### miRNA data exploration of uEVs isolated from AUD patients

Before assessing the miRNA profile of uEVs in healthy individuals and AUD patients, we characterized the isolated uEVs using TEM, Western blot, and NTA (Fig. 2). TEM imaging confirmed that the nanoscale particles displayed the expected exosomal morphology and size (∼100 nm in diameter; Fig. 2A). These uEVs also expressed the exosomal markers CD63, CD9, and CD81 (tetraspanins) and showed no evidence of cytosolic contamination, as indicated by the lack of calnexin detection (Fig. 2B). NTA analysis further supported these observations, revealing a predominant particle size within the 100–200 nm range, consistent with EVs characteristics (Fig. 2C). Following the characterization of uEVs, an exploratory analysis of the miRNA profiles across all samples was performed. The resulting correlation heatmap revealed that most AUD samples clustered separately from controls, with partial segregation by sex (Fig. 2D).

**Figure 2.**
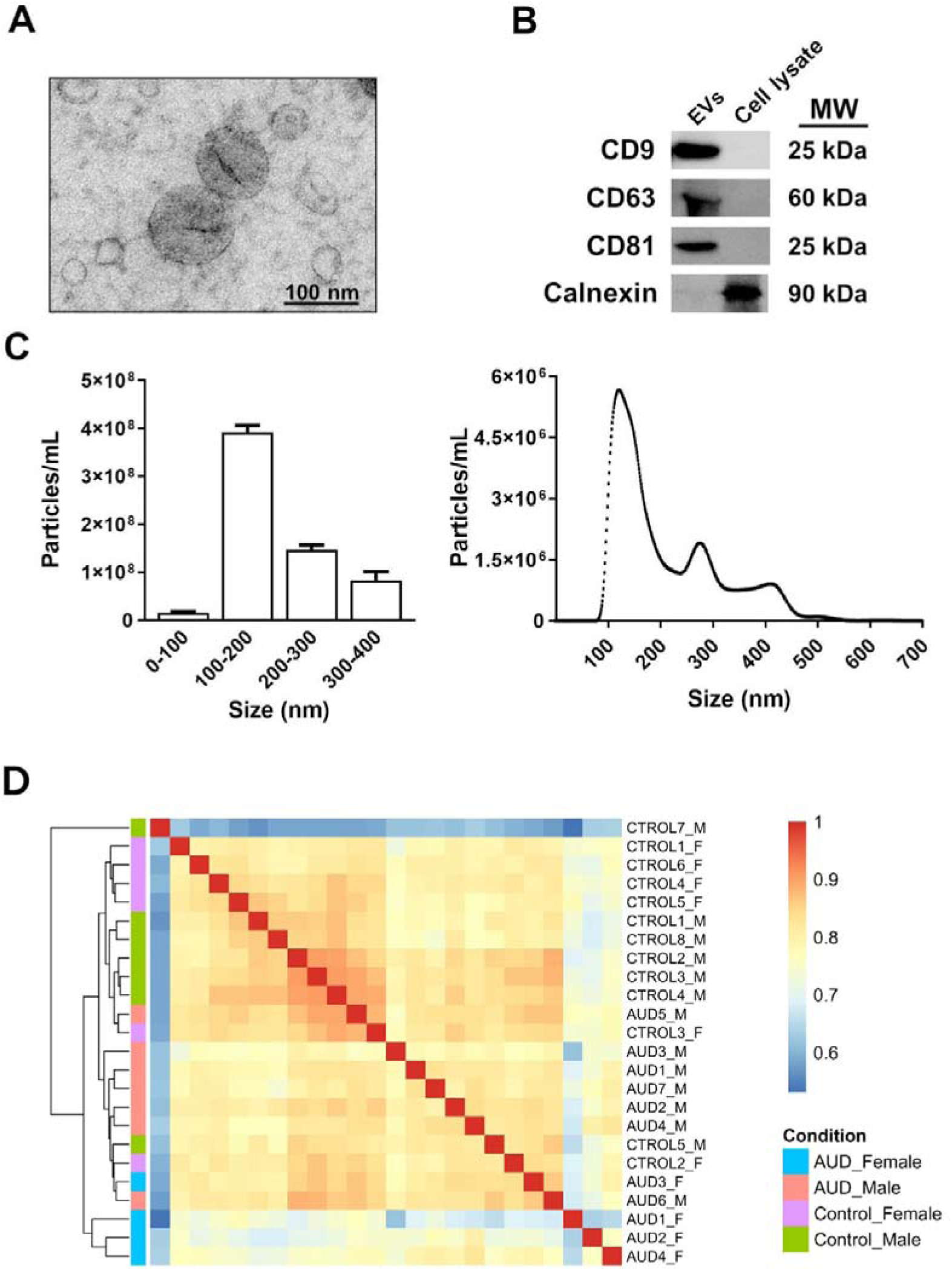
Characterization of uEVs. **A)** Transmission electron microscopy (TEM) image of human uEVs following isolation. **B)** Immunoblot detection of canonical EV markers (CD9, CD63, and CD81) in uEVs, alongside astroglial cell lysates used as a positive control for calnexin. Assessment of calnexin expression was performed to confirm the absence of cytosolic protein contamination in the EV preparations. Representative blots for each protein are presented. **C)** Nanoparticle tracking analysis (NTA) of uEVs, illustrating particle size distribution and particle concentration. **D)** Correlation heatmap illustrating sample-to-sample similarity based on miRNA expression profiles. Warmer colors indicate higher correlation, revealing the distribution and clustering patterns among samples.

### Analysis of miRNA expression profile of AUD patient uEVs

To explore potential biomarkers in uEVs from AUD patients, we analyzed samples from both female and male AUD individuals as well as healthy controls. The study focused on the role of miRNAs across the four experimental groups, defined by sex and alcohol consumption status, using miRNA-sequencing data obtained from corresponding uEV samples. Differential expression analysis with DESeq2 identified miRNAs that were significantly altered between AUD and control groups within each sex (IF and IM comparisons). Figure 3A–B shows the upregulated (LFC > 0) and downregulated (LFC < 0) miRNAs, with the full dataset available on the web platform (https://bioinfouvcipf.shinyapps.io/miRNAuEVs-AUD/). In females, twelve miRNAs were significantly upregulated and two downregulated, whereas in males, three miRNAs were upregulated and three downregulated. These data indicate a higher number of upregulated miRNAs in females compared to males, suggesting sex-specific responses to chronic alcohol exposure. In addition, we also detected four significantly altered miRNAs in the IS comparison, two upregulated and two downregulated miRNAs (Table S2 and web platform - study overview [https://bioinfouvcipf.shinyapps.io/miRNAuEVs-AUD/]). The Upset plot (Fig. 3B) further highlighted that one miRNA (hsa-miR-4787-5p) was commonly upregulated in both IF and IM comparisons, suggesting a shared response to alcohol exposure across sexes.

**Figure 3.**
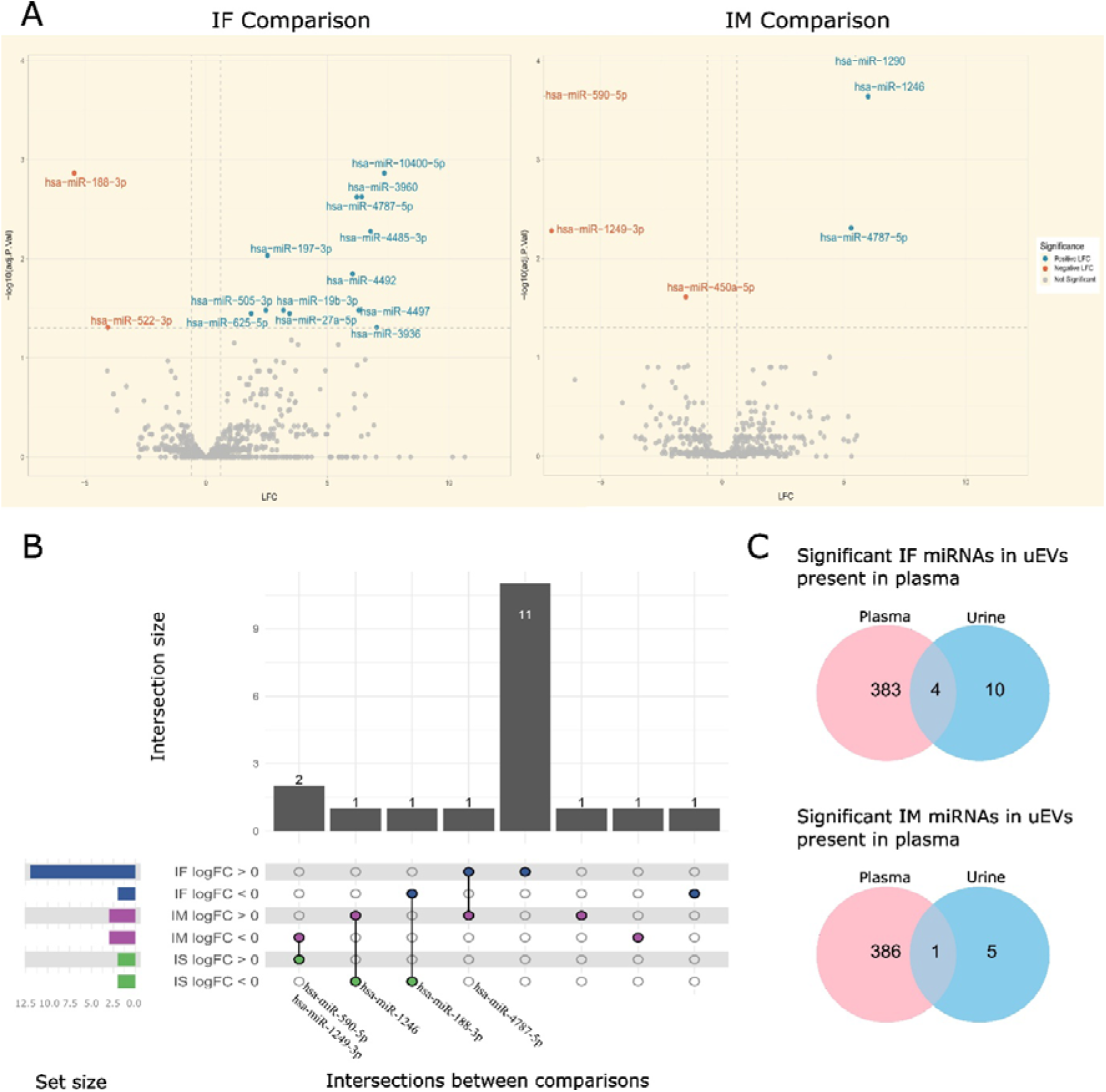
Differential miRNA expression in uEVs from AUD patients versus controls. **A)** Volcano plots of all miRNAs in the IF (AUD Impact in Females) and IM (AUD Impact in Males) comparisons (colored by direction of change: miRNAs with a positive log fold change (LFC > 0) display upregulation (blue dots) and miRNAs with a negative LFC (LFC < 0) display downregulation (red dots) in AUD patients (adjusted p ≤ 0.05). **B)** Upset plot showing shared and unique significantly altered miRNAs across IF, IM, and IS comparisons. miRNAs labeled under the dots correspond to the respective intersections. Intersection plot displaying shared and unique significantly altered miRNAs across IF, IM, and IS comparisons. miRNAs listed below each dot are present in the comparisons corresponding to that intersection. **C)** Venn diagrams showing overlap of significantly altered miRNAs in uEVs (adjusted p ≤ 0.05) in both comparisons (IF and IM), that were also detected in plasma EVs.

Then, Figure 3C compared the uEV miRNome data with previously published plasma EV miRNome data from male and female AUD patients and controls [22]. Technical differences in EV isolation and miRNA extraction between urine and plasma samples are summarized in Table S3. Compared with blood, urine offers several key advantages, including non-invasive collection, larger sample volumes, and reduced proteomic complexity [43], making it a particularly attractive biofluid for omics-based investigations. Venn diagram analyses showed that four miRNAs (hsa-miR-197-3p, hsa-miR-19b-3p, hsa-miR-505-3p, hsa-miR-625-5p) with significant changes in uEVs were also detected in plasma EVs for the IF comparison, while one miRNA (hsa-miR-450a-5p) overlapped in the IM comparison.

### Functional analysis of miRNA of AUD patient uEVs: KEGG and Reactome pathways

To investigate the biological functions associated with miRNAs present in uEVs from AUD patients, we performed a Gene Set Enrichment Analysis (GSEA) on experimentally validated target genes of the differentially expressed miRNAs using the KEGG and Reactome pathway databases. Differentially expressed miRNAs identified in the two independent comparisons—AUD females versus female controls (IF) and AUD males versus male controls (IM)—were used as input for the analysis. This approach revealed multiple significantly enriched functional pathways across both sex-specific comparisons, highlighting both shared and distinct biological signatures associated with AUD.

KEGG pathway enrichment analysis revealed broad functional categories related to vesicle trafficking and endocytosis, signal transduction and kinase activity, cell cycle regulation and apoptotic processes, immune and inflammatory responses, and cancer-related proliferation pathways, among others (Fig. 4A). Most enriched KEGG terms displayed negative log odds ratio (LOR) values in both IF and IM comparisons, indicating a predominant association with gene sets derived from downregulated miRNAs. Despite these shared features, some sex-dependent patterns were observed. Pathways associated with cell adhesion and cytoskeletal dynamics were detected exclusively in IF, whereas vesicle trafficking/endocytic and immune–inflammatory pathways were mainly represented in IM. In addition, cancer- and proliferation-related pathways, together with protein/RNA processing and signal transduction pathways, were more abundant in IF, while cell cycle and apoptotic pathways were shared between both comparisons.

**Figure 4.**
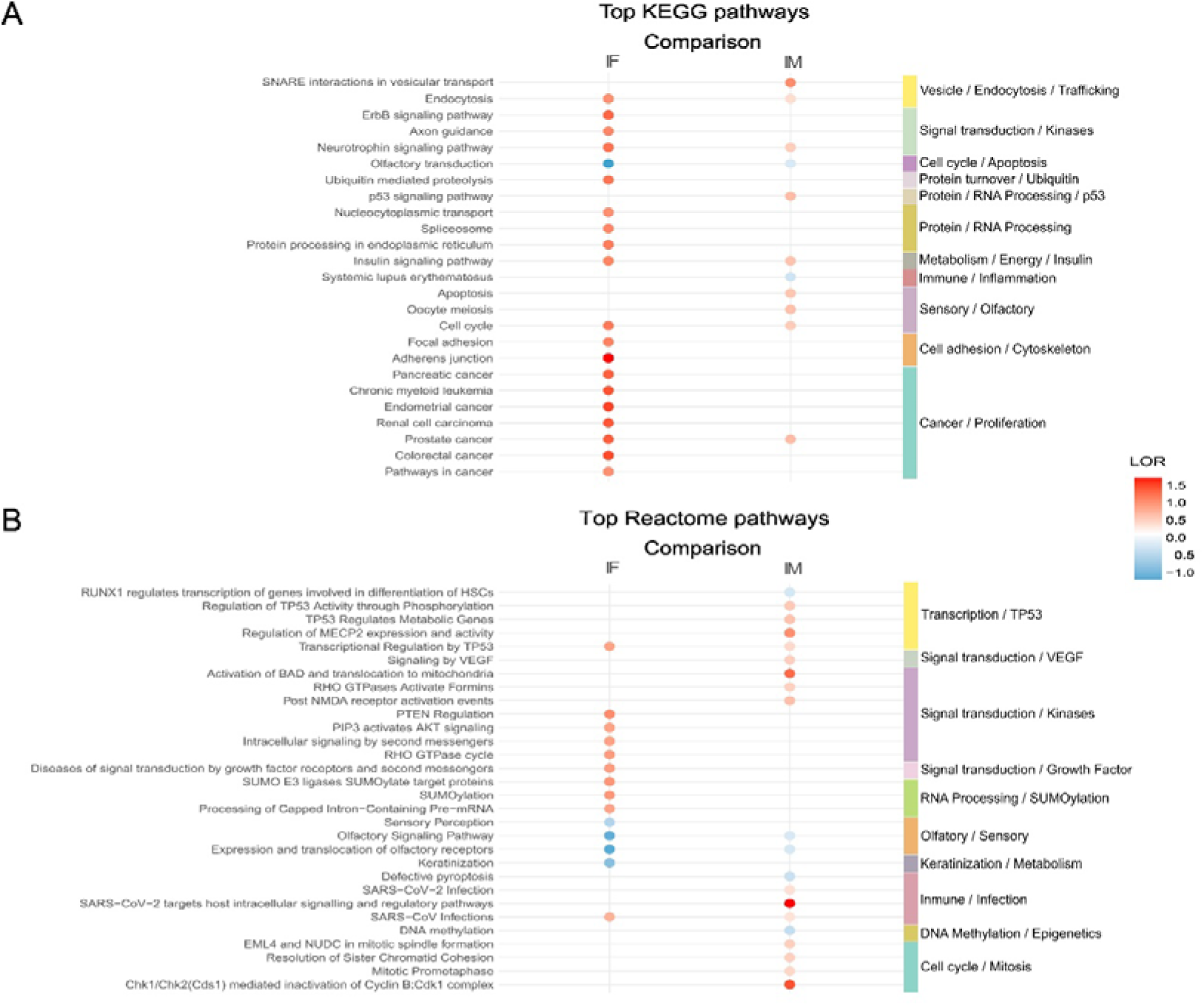
Functional enrichment of miRNAs in AUD patient uEVs: top KEGG and Reactome pathways in IF and IM. **A)** Dot plot showing the top 20 significant Kyoto Encyclopedia of Genes and Genomes (KEGG) terms for the IF and IM comparisons (ordered by adjusted p-value, lowest to highest). The y-axis represents the KEGG terms, and the x-axis shows the two comparisons: IF (AUD Impact in Females) and IM (AUD Impact in Males). Dot color indicates the log odds ratio (LOR) value and the colors shown in the right column indicate general functional categories. **B)** Dot plot displaying the top 20 significant Reactome pathways for the IF and IM comparisons, with the same layout and visual conventions as in (A).

Reactome pathway analysis further supported the presence of shared and sex-specific functional signatures (Fig. 4B). Enriched Reactome pathways were predominantly associated with TP53-mediated transcription, VEGF and growth factor signaling, kinase-driven signal transduction, immune- and infection-related processes, and cell cycle and mitotic progression, among others. TP53-related, VEGF signaling, DNA methylation/epigenetic, and mitotic cell cycle pathways were identified predominantly or exclusively in IM, whereas growth factor signaling, RNA processing/SUMOylation, and keratinization and metabolic pathways were specific to IF. Kinase-mediated signaling pathways were common to both comparisons, while immune- and infection-related pathways were more extensively represented in IM.

### Gene Ontology–based functional clustering of miRNA target genes

We next performed a Gene Ontology (GO)–based functional clustering to characterize the collective biological role of the miRNA target genes. This semantic similarity analysis focused on the Biological Processes (BP) and Cellular Components (CC) categories, as they provide the most interpretable insights into system-level biology. The enriched terms within these categories were grouped into non-redundant clusters, revealing the principal functional modules potentially under miRNA regulation in AUD. Figure 5 shows that GO terms associated with target genes of upregulated miRNAs are shown in blue, whereas those linked to downregulated miRNAs are shown in red; BP and CC categories are indicated by orange and green labels, respectively, and word clouds highlight representative terms within each functional cluster.

**Figure 5.**
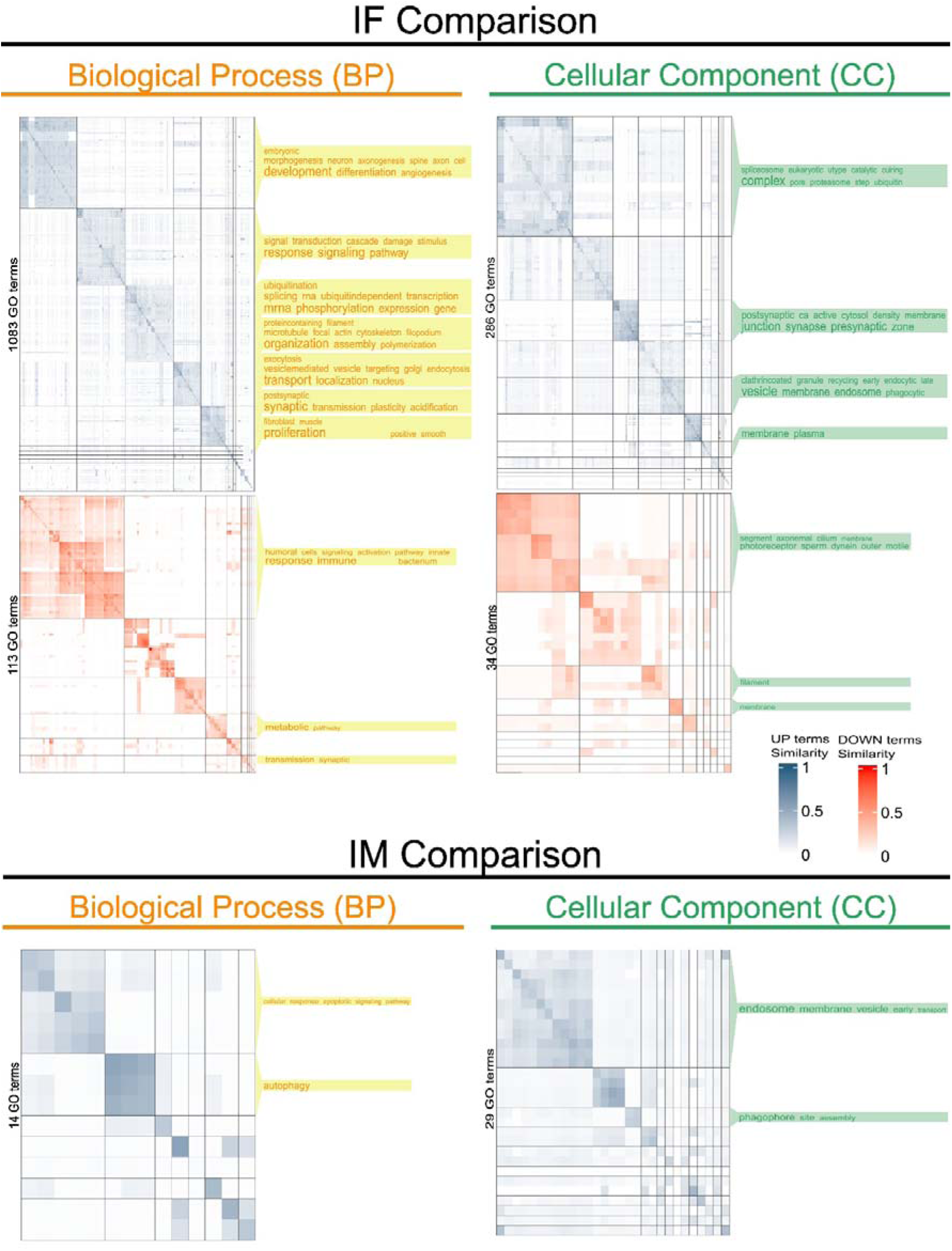
Gene set analysis of GO Biological Process and Cellular Component terms for IF and IM comparisons, clustered by semantic similarity. Clustering of significant Gene Ontology (GO) Biological Process (BP) (purple labels) and Cellular Component (CC) (green labels) terms for both IF and IM comparisons based on semantic similarity. Word clouds summarize representative terms for each cluster. Red clusters correspond to GO terms associated with target genes of downregulated miRNAs, whereas blue clusters represent those linked to upregulated miRNAs.

In the IF comparison (AUD females vs. controls), BP terms associated with upregulated miRNAs segregated into multiple functional clusters reflecting developmental and differentiation-related processes, angiogenesis and neuronal morphogenesis, cellular responses and signaling pathways, regulation of gene expression and mRNA processing, ubiquitination, cytoskeletal organization and cellular assembly, vesicle-mediated trafficking, synaptic function and plasticity, and cellular proliferation. In contrast, BP terms associated with downregulated miRNAs in IF were primarily linked to innate and humoral immune responses, metabolic processes, and synaptic transmission. At the CC level, upregulated miRNA–associated terms clustered around spliceosome-and ubiquitin-related complexes, pre- and postsynaptic compartments, endosomal vesicles, and plasma membrane structures, whereas downregulated miRNA–associated CC terms were mainly related to ciliary structures, filament organization, and specialized membrane components.

In the IM comparison (AUD males vs. controls), BP terms associated with upregulated miRNAs were predominantly enriched in cellular response pathways, apoptotic signaling, and autophagy-related processes. Correspondingly, CC terms were mainly associated with endosomal compartments, transport vesicles, and phagophore assembly sites. GO BP and CC terms linked to downregulated miRNAs were not represented in IM due to an insufficient number of significant terms for robust enrichment analysis. A summary of the number of significant GSA results for each comparison is provided in Table S4, while Figures S2, S3 and S4 illustrate the overlap of the top 20 significant GO BP, CC and MF terms between IF and IM, respectively.

### Web platform

Finally, we developed the interactive web platform “miRNAuEVs-AUD” (https://bioinfouvcipf.shinyapps.io/miRNAuEVs-AUD/) that allows users to explore the miRNA profiles presented in this study in a dynamic and user-friendly way. The platform provides access to uEV miRNA datasets and differential expression results from all study comparisons (IF, IM, and IS), enabling users to examine both sex-specific differences and alcohol-related changes. Users can query specific miRNAs, adjust analysis parameters, and generate custom visualizations, expanding the information available in the manuscript on AUD-associated miRNA biomarkers.

## Discussion

We have previously reported an AUD-sex signature (e.g., hsa-miR-1301-3p, PC39:4) revealing sex-divergent biological responses to alcohol and demonstrating the value of multi-omic integration. In the individual miRNome approach, we uncover opposing functional alterations between sexes in immunity, oxidative stress, and autophagy pathways [22]. Extending these findings, this study pioneers the systematic profiling of miRNAs within uEVs from a clinically characterized AUD cohort, establishing a novel molecular signature from an accessible, non-invasive biofluid. While uEVs have traditionally been associated with renal and urinary tract disorders [12], growing evidence now supports their relevance in cancer [44] and neurological conditions [45–47]. By incorporating sex-stratified analyses, we aim to identify potential biomarkers for the diagnosis and prognosis of chronic heavy alcohol consumption, positioning uEVs as dynamic and readily accessible tools for disease monitoring [44, 47, 48].

Differential analysis of the miRNome revealed pronounced sexual dimorphism in uEV miRNA signatures, with distinct molecular responses to AUD in females and males. While some miRNAs were consistently dysregulated across sexes, others showed clear sex-specific regulation, reflecting partially divergent regulatory mechanisms. These findings align with our previous multi-omic analysis in plasma EVs, which revealed sex-divergent biological responses to alcohol and underscored the value of integrating lipidomic and miRNomic data [22]. Comparison with plasma-derived EVs further suggests that uEVs capture complementary molecular information not fully represented in systemic circulation, consistent with previous studies showing that uEVs contain a larger fraction of tissue-derived molecular signals compared to plasma EVs [48].

Chronic alcohol consumption is closely associated with progressive liver injury, ranging from steatosis to cirrhosis, processes in which EV-mediated intercellular communication plays a key role [49, 50]. Since the AUD patients in our cohort had been actively drinking in the weeks leading up to urine collection, they offer insight into alcohol-related metabolic changes, as opposed to the consequences of end-stage organ pathology. However, the increased levels of hsa-miR-1290 and hsa-miR-197-3p observed in the IM and IF comparisons, respectively, point to a direct involvement of these miRNAs in alcohol-related liver injury [50–52]. hsa-miR-1290 has been described in cirrhosis and non-malignant chronic diseases as a regulator of cellular homeostasis and autophagy, whereas hsa-miR-197-3p is functionally linked to the IL-6/STAT3 axis, a central pathway in hepatic inflammation and alcohol-induced damage [51, 52]. Beyond the liver, both miRNAs participate in the modulation of systemic inflammatory responses [51–54]. In this context, the downregulation of hsa-miR-522-3p in IF is particularly relevant, given its role in regulating phagocytosis and inflammation resolution through the HMGB1–PTGR1 axis [55], suggesting impaired anti-inflammatory mechanisms in women with AUD. Moreover, the effects of alcohol on the central nervous system are also reflected in the uEV signature, where the upregulation of hsa-miR-19b-3p in IF and hsa-miR-1246 in IM has been associated with neuroinflammation, neuronal stress, dysregulation of PI3K/AKT/mTOR signaling, and altered synaptic signaling [56–58], processes widely implicated in AUD-related neurological disorders. Consistent with these findings, functional enrichment analysis of differentially expressed miRNAs revealed significant overrepresentation of pathways related to kinase-mediated signaling, synaptic function and plasticity, immune–inflammatory processes, autophagy, and vesicle trafficking/endocytosis, reinforcing the involvement of these regulatory networks in liver injury, systemic inflammation, and neuronal dysfunction associated with AUD.

A second major axis of our findings links the uEV miRNome to canonical processes of carcinogenesis and cellular plasticity, consistent with evidence that microRNAs mediate the effects of environmental risk factors on cancer progression [59]. In IM, the upregulation of hsa-miR-1290 and hsa-miR-1246, and in IF the differential detection of hsa-miR-27a-5p, is particularly noteworthy, as these miRNAs have been widely described as regulators of proliferation, invasiveness, and therapy resistance across multiple cancer types (e.g., colorectal, breast, and lung cancer) [58, 60–63]. However, the literature also attributes context-dependent tumor-suppressive functions to some of these miRNAs. For instance, hsa-miR-27a-5p has been described as a tumor suppressor in several models, modulating pathways such as PI3K/AKT and MAPK [62, 64]. This duality suggests that its presence in uEVs from patients with AUD may reflect not only pro-oncogenic processes but also compensatory responses to chronic alcohol-induced tissue damage. In agreement, KEGG and Reactome enrichment analyses revealed significant overrepresentation of pathways related to growth factor signaling, kinase-driven signal transduction, cell cycle regulation, mitotic processes, and cancer-associated pathways, supporting the notion that the uEV miRNA signature in AUD reflects not only chronic inflammation but also a reprogramming of proliferation and cellular plasticity circuits linked to carcinogenic risk.

From a translational perspective, several of the identified miRNAs further support the potential of EVs as a source of organ-specific and systemic biomarkers in AUD [50]. The downregulation of hsa-miR-590-5p in IM, previously associated with the regulation of VEGF signaling, angiogenesis, and oxidative stress, may reflect alterations in endothelial and microvascular damage control induced by alcohol [65]. In IF, the upregulation of hsa-miR-505-3p and hsa-miR-3960, both previously described as biomarkers in liver diseases and other pathological contexts, supports the existence of a uEV signature with potential diagnostic and prognostic value [66–68]. Additionally, the increased levels of hsa-miR-4492 in IF are particularly noteworthy, as this miRNA has been described as a central node in multiple lncRNA–miRNA–mRNA regulatory axes [68–70], suggesting that its dysregulation may contribute to a broader reorganization of gene regulatory circuits in response to AUD. Finally, the exclusive detection of hsa-miR-4787-5p in uEVs but not in plasma-derived EVs, together with its prior association with renal processes, supports a urinary-compartment-associated signal and warrants further validation and additional investigation of its potential utility as a specific biomarker of subclinical renal injury in patients with AUD [71–73]. These results also highlight the distinctive value of uEVs as a biological window into alcohol-affected target organs.

AUD exerts complex biological effects, increasing the risk of cardiovascular, neurological, metabolic, hepatic, and oncological diseases [74, 75]. Although sex-based differences in AUD, particularly in inflammatory responses, have been reported [76, 77], they are often underestimated in research. Historically, the underrepresentation of females in AUD studies has generated critical gaps in knowledge, diagnosis, and treatment, highlighting the need for more sex-inclusive and sensitive approaches [78]. In this context, our study employed an innovative strategy—profiling the uEV miRNome—to explore these sex-specific molecular signatures. However, this strategy, while promising, is subject to important limitations. First, the total number of participants, including AUD males, was relatively small due to the high comorbidity and access barriers characteristic of this clinical population, a factor that may affect the generalizability of the findings. Second, the recruitment of AUD females was particularly challenging owing to sociocultural factors such as underreporting and stigma, resulting in a reduced sample size and limited statistical power for sex-stratified comparisons [79]. Finally, the absence of a consensus on optimal miRNA normalization methods for EV studies hinders direct comparisons with other datasets. Moving forward, it will be essential to functionally validate the candidate miRNAs identified here in relevant experimental models and to confirm these preliminary findings in larger, independent cohorts with balanced sex representation.

In conclusion, our findings demonstrate that uEVs capture complex and sex-dependent molecular responses to chronic alcohol consumption, providing novel insights into the regulatory networks underlying AUD pathophysiology. The uEV miRNome revealed distinct signatures associated with liver injury, systemic inflammation, neurobiological alterations, cellular plasticity, and carcinogenesis-related pathways, many of which were differentially represented between females and males. Collectively, these results underscore the value of uEVs as a non-invasive and sensitive biofluid for identifying sex-specific and organ-related molecular alterations associated with chronic alcohol exposure, supporting their potential utility as biomarkers for AUD diagnosis, prognosis, and systemic monitoring.

## Declarations

### Ethics approval and consent to participate

Human urine samples were used in accordance with the Declaration of Helsinki and were approved by the Ethics Committee of the University Hospital of Salamanca (PI 2023 07 1389), and written informed consent was obtained from each participant.

### Supporting information

Table S1. Clinical and demographic characteristics of participants with chronic alcohol consumption.

Table S2. Differentially expressed miRNAs in uEVs for IF, IM, and IS comparisons

Table S3. Technical differences in EV isolation and miRNA analysis between urine and plasma samples.

Table S4. Number of significant GO terms identified by GSEA for each comparison.

Figure S1. Western blots of CD9, CD63, CD81, and calnexin.

Figure S2. Dot plot showing the top 20 significant BP terms for the IF and IM comparisons.

Figure S3. Dot plot showing the top 20 significant CC terms for the IF and IM comparisons.

Figure S4. Dot plot showing the top 20 significant MF terms for the IF and IM comparisons.

### Data availability statement

Datasets and analysis scripts supporting the findings of this study are available through Github, https://github.com/Blancamartinurdiales/Urinary-extracellular-vesicles-reveal-a-sex-specific-miRNome-profile-in-AUD-patients, and in a web platform: https://bioinfouvcipf.shinyapps.io/miRNAuEVs-AUD/

### Competing interests

The authors declare that they have no competing interests.

### Funding

This work has been supported by grants from the Spanish Ministry of HealthLPNSD (2023-I024), GVA (CIAICO/2024/122 and CIAICO/2023/149), the Primary Addiction Care Research Network (RD21/0009/0005 and RD24/0003/0017), FEDER Funds, GVA and the Instituto de Salud Carlos III (ISCIII) through the project PI20/00743 and PI25/00279, co-funded by the European Union and the Junta de Castilla y León (GRS 2916/A1/2023 and GRS3005/A1/2024), PID2021-124430OA-I00 and PID2023-146865OB-I00 funded by MCIN/AEI/10.13039/501100011033/FEDER, UE (“A way to make Europe”). MRP holds a Sara Borrell contract (CD22/00054) funded by Instituto de Salud Carlos III (ISCIII) and cofunded by the European Union–Next Generation EU.

### Authors contributions

BMU analyzed the data; MP, CPC and FGG designed and supervised the bioinformatics analysis; MLAS, MRP, DPM, and MM obtained human urine samples and collected clinical data; SM isolated EVs from human urine and miRNAs; BMU designed and implemented the web tool; BMU and MP wrote the manuscript; MP and BMU designed the graphical abstract; BMU and MP interpreted the results; BMU, CPC, MM, MP and FGG writing-review and editing; MP conceived the work. All authors have read and approved the final manuscript.

## Acknowledgments

The authors thank the Principe Felipe Research Center (CIPF) for providing access to the cluster, co-funded by European Regional Development Funds (FEDER) in the Valencian Community 2014-2020. The authors also thank the Genomics and Epigenetics section of the Central Research Unit of Medicine at the University of Valencia.

## Notes

https://bioinfouvcipf.shinyapps.io/miRNAuEVs-AUD/

### Competing Interest Statement

The authors have declared no competing interest.

