## Supplementary material for "Urinary extracellular vesicles reveal a sex-specific miRNome profile in alcohol use disorder patients"

**Table S1.** Characteristics of study individuals displaying chronic alcohol consumption.

| **PARAMETER** | **Men (n=7)** | **Reference values** | **Women (n=4)** | **Reference values** |
| --- | --- | --- | --- | --- |
| **Age (years)** | 49.1 (13.1) |  | 47.8 (7.4) |  |
| **Total bilirubin (mg/dL)** | 0.9 (0.5) | 0.2-1.1 | 0.5 (0.2) | 0.2-0.9 |
| **AST (U/L)** | 68.0 (30.5) | <34 | 59.0 | <31 |
| **ALT (U/L)** | 50.6 (23.9) | 10-49 | 17.8 (6.6) | 10-49 |
| **FA (U/L)** | 77.7 (37.7) | 46-116 | 92.8 (25.8) | 46-116 |
| **LDH (U/L)** | 179.3 (19.6) | 120-246 | 220.0 (29.7) | 120-246 |
| **GGT (U/L)** | 196.2 (229.2) | <73 | 53.3 (36.0) | <38 |
| **Proteins (g/dL)** | 7.1 (0.6) | 5.7-8.2 | 6.7 (0.5) | 5.7-8.2 |
| **Albumin (g/dL)** | 4.4 (0.2) | 3.2-4.8 | 4.3 (0.3) | 3.2-4.8 |
| **Ferritin (ng/mL)** | 249.4 (143.9) | 15-200 | 67.8 (39.3) | 15-200 |
| **Hemoglobin (g/dL)** | 15.9 (1.4) | 13-17 | 14.3 (0.6) | 12-15 |
| **Hematocrit (%)** | 43.5 (13.2) | 40-50 | 43.2 (3.2) | 25-45 |
| **MCV (fL)** | 94.5 (7.8) | 80-98 | 94.8 (3.7) | 80-98 |
| **MCH (pg)** | 31.6 (2.42) | 27-32 | 32.0 (1.6) | 27-32 |
| **Leukocytes (x10^3^ cel/μL)** | 7.6 (1.2) | 3.8-11 | 9.2 (4.31) | 3.8-11 |
| **Neutrophils (x10^3^ cel/μL)** | 4.5 (1.41) | 1.8-7 | 5.6 (3.5) | 1.8-7 |
| **Lymphocytes (x10^3^ cel/μL)** | 2.1 (0.9) | 1-4.5 | 2.5 (0.6) | 1-4.5 |
| **Platelets (x10^3^ cel/μL)** | 239.0 (58,1) | 140-450 | 203.0 (83.7) | 140-450 |
| **Total cholesterol (mg/dL)** | 173.1 (46.9) | <200 | 192.8 (39.7) | <200 |
| **Triglycerides (mg/dL)** | 138.9 (100.6) | <150 | 351.3 (487.6) | <150 |
| **Prothrombin activity (%)** | 84.5 (19.1) | 70-120 | 110.9 (61.7) | 70-120 |
| **APTT (segundos)** | 23.6 (12.9) | 21.6-30 | 26.5 (0.6) | 21.6-30 |
| **Fibrinogen (mg/dL)** | ND | 170-420 | 433.7 | 130-400 |
| **D-dimer (μg/mL)** | ND | 0-0.6 | 0.6 | 0-0.8 |

Note: mean (standard deviation). AST: aspartate aminotransferase. ALT: alanine aminotransferase. AP: alkaline phosphatase. LDH: lactate dehydrogenase. GGT: gamma glutamyl transferase. MCV: mean corpuscular volume. MCH: mean corpuscular hemoglobin. APTT: activated partial thromboplastin time.

### Table S2. Differentially expressed miRNAs in uEVs for IF, IM, and IS comparisons.

|  | **miRNA** | **adjusted p** | **LFC** |  |
| --- | --- | --- | --- | --- |
| IF | hsa-miR-10400-5p | 0.001364 | 7.351466 | Upregulated |
|  | hsa-miR-3960 | 0.002364 | 6.416949 |  |
|  | hsa-miR-4787-5p | 0.002377 | 6.214586 |  |
|  | hsa-miR-4485-3p | 0.005283 | 6.776318 |  |
|  | hsa-miR-197-3p | 0.00928 | 2.533609 |  |
|  | hsa-miR-4492 | 0.01423 | 6.042533 |  |
|  | hsa-miR-19b-3p | 0.033111 | 3.190401 |  |
|  | hsa-miR-505-3p | 0.033111 | 2.46366 |  |
|  | hsa-miR-4497 | 0.033111 | 6.2973 |  |
|  | hsa-miR-27a-5p | 0.035587 | 3.442897 |  |
|  | hsa-miR-625-5p | 0.035587 | 1.860727 |  |
|  | hsa-miR-3936 | 0.049177 | 7.035273 |  |
|  | hsa-miR-188-3p | 0.001364 | -5.43538 | Downregulated |
|  | hsa-miR-522-3p | 0.049177 | -4.05139 |  |
| IM | hsa-miR-1290 | 4.03E-05 | 5.4646349 | Upregulated |
|  | hsa-miR-1246 | 0.00023 | 5.9975218 |  |
|  | hsa-miR-4787-5p | 0.004927 | 5.2961555 |  |
|  | hsa-miR-590-5p | 0.00023 | -18.708080 | Downregulated |
|  | hsa-miR-1249-3p | 0.005249 | -7.0088339 |  |
|  | hsa-miR-450a-5p | 0.024462 | -1.4918456 |  |
| IS | hsa-miR-1249-3p | 0.04543 | 9.648869 | Upregulated |
|  | hsa-miR-590-5p | 0.04543 | 21.62832 |  |
|  | hsa-miR-1246 | 0.04543 | -6.99222 | Downregulated |
|  | hsa-miR-188-3p | 0.04543 | -5.37768 |  |

### Table S3. Technical differences in EV isolation and miRNA analysis between urine and plasma samples.

| Sample type | **Urine** | **Plasma** |
| --- | --- | --- |
| AUD group | 7 males and 4 females | 6 males and 3 females |
| Control group | 7 males and 6 females | 5 males and 6 females |
| Starting sample volume | 21 mL/sample | 250 μL/sample |
| EVs isolation method | Deferential ultracentrifugation | Total exosome isolation kit |
| EVs characterization | NTA, TEM, Western blot | NTA, TEM, Western blot |
| miRNA extraction | Total Exosome RNA Isolation Kit (Invitrogen) | Total Exosome RNA Isolation Kit (Invitrogen) |
| Small RNA library preparation | NEXTFLEX® Small RNA-Seq Kit v4 with UDIs | TruSeq Small RNA Library Preparation Kit |
| Indexing strategy | Unique dual indexes (UDIs) | Single indexing |
| Library quality control | Bioanalyzer 2100 (High Sensitivity DNA chips) | TapeStation (High Sensitivity D1000 ScreenTape) |
| Sequencing platform | NextSeq 550 | MiSeq™ |
| Total miRNAs | 713 | 387 |

### Table S4. Number of significant GO terms identified by GSEA for each comparison.

| **Comparation** | **LFC** | **Nº of significant GSA results** | **GO ontology** |
| --- | --- | --- | --- |
| IF | UP | 1083 | Biological Process  BP |
|  | DOWN | 113 |  |
|  | UP | 286 | Cellular Component CC |
|  | DOWN | 34 |  |
| IM | UP | 14 | Biological Process  BP |
|  | DOWN | 2 |  |
|  | UP | 26 | Cellular Component CC |
|  | DOWN | 6 |  |

**Figure S1.** Western blots of CD9, CD63, CD81, and calnexin are shown.

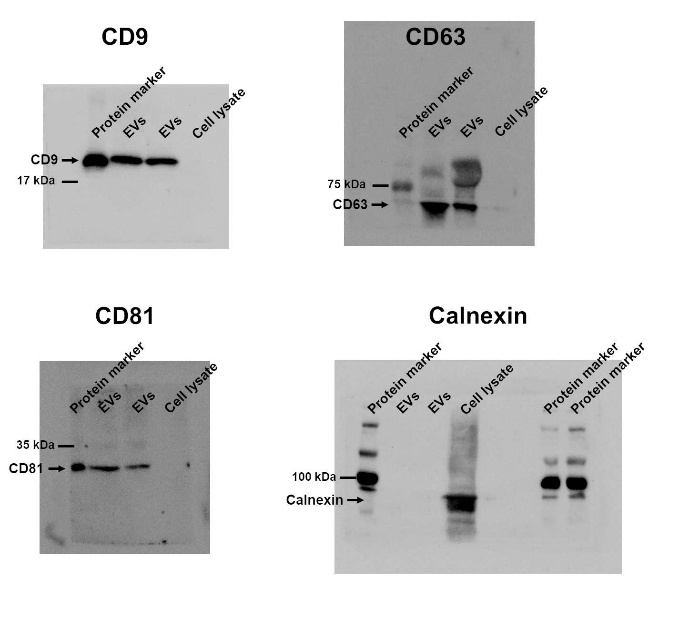

### Figure S2. Dot plot showing the top 20 significant BP terms for the IF and IM comparisons (ordered by adjusted p-value, lowest to highest).

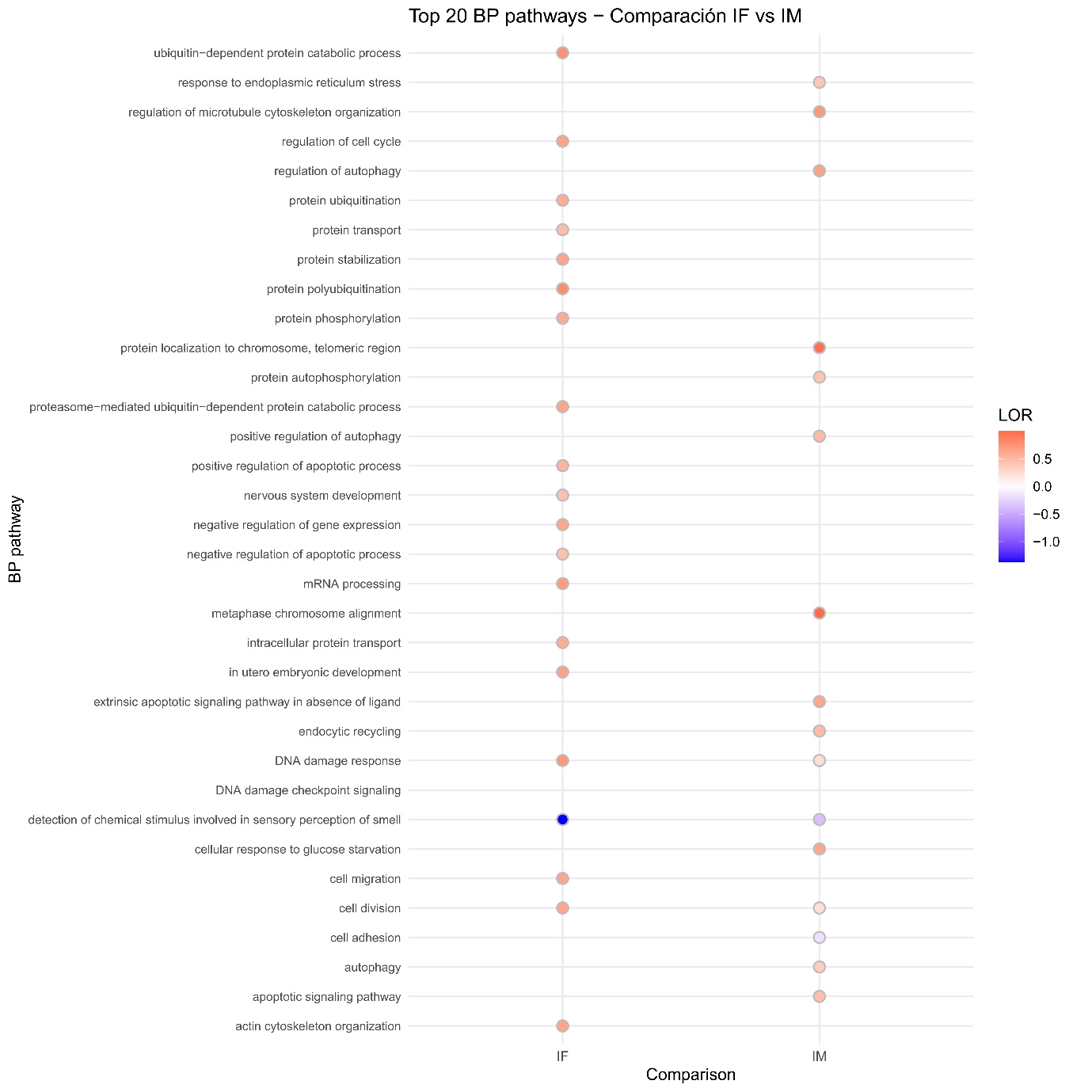

### Figure S3. Dot plot showing the top 20 significant CC terms for the IF and IM comparisons (ordered by adjusted p-value, lowest to highest).

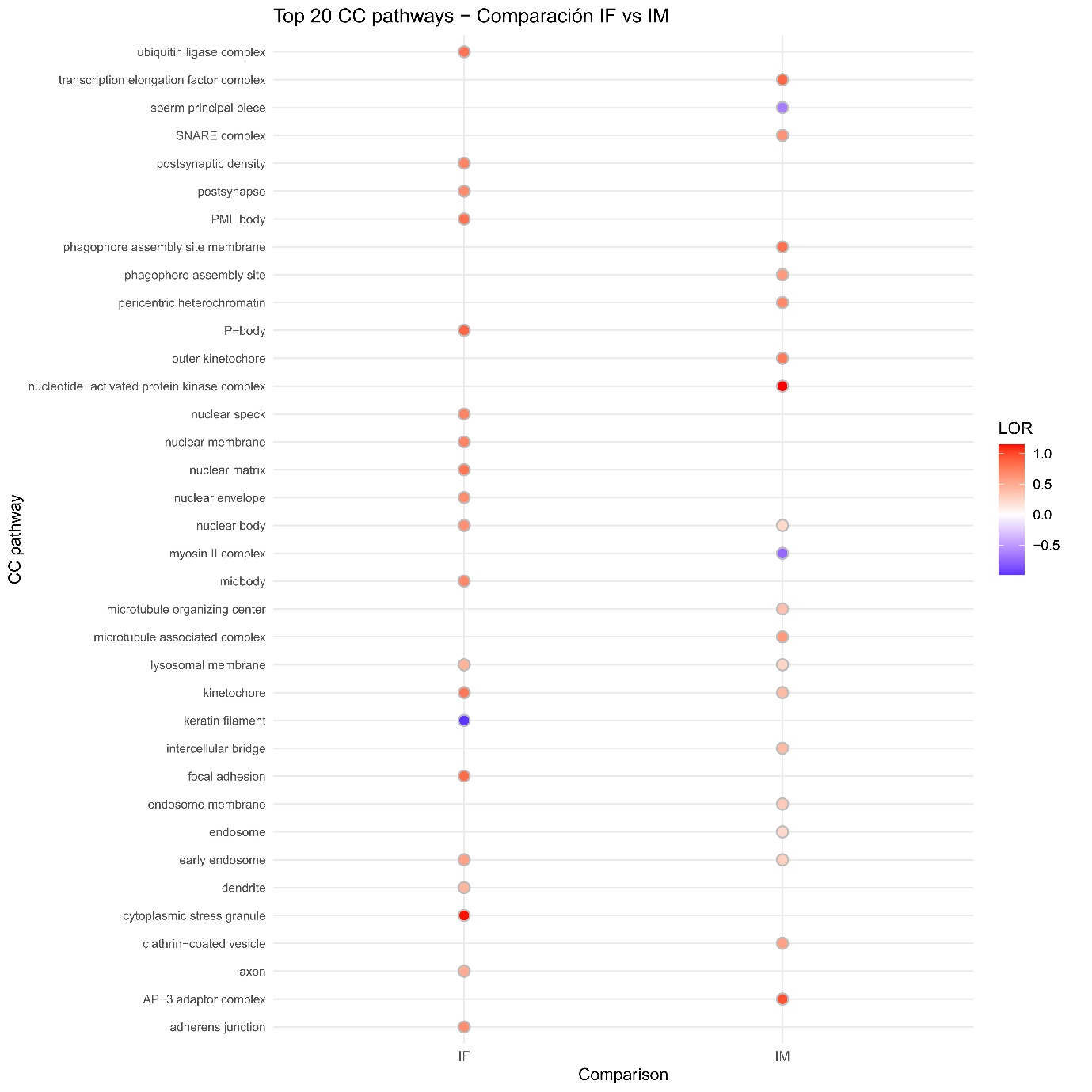

### Figure S4. Dot plot showing the top 20 significant MF terms for the IF and IM comparisons (ordered by adjusted p-value, lowest to highest).

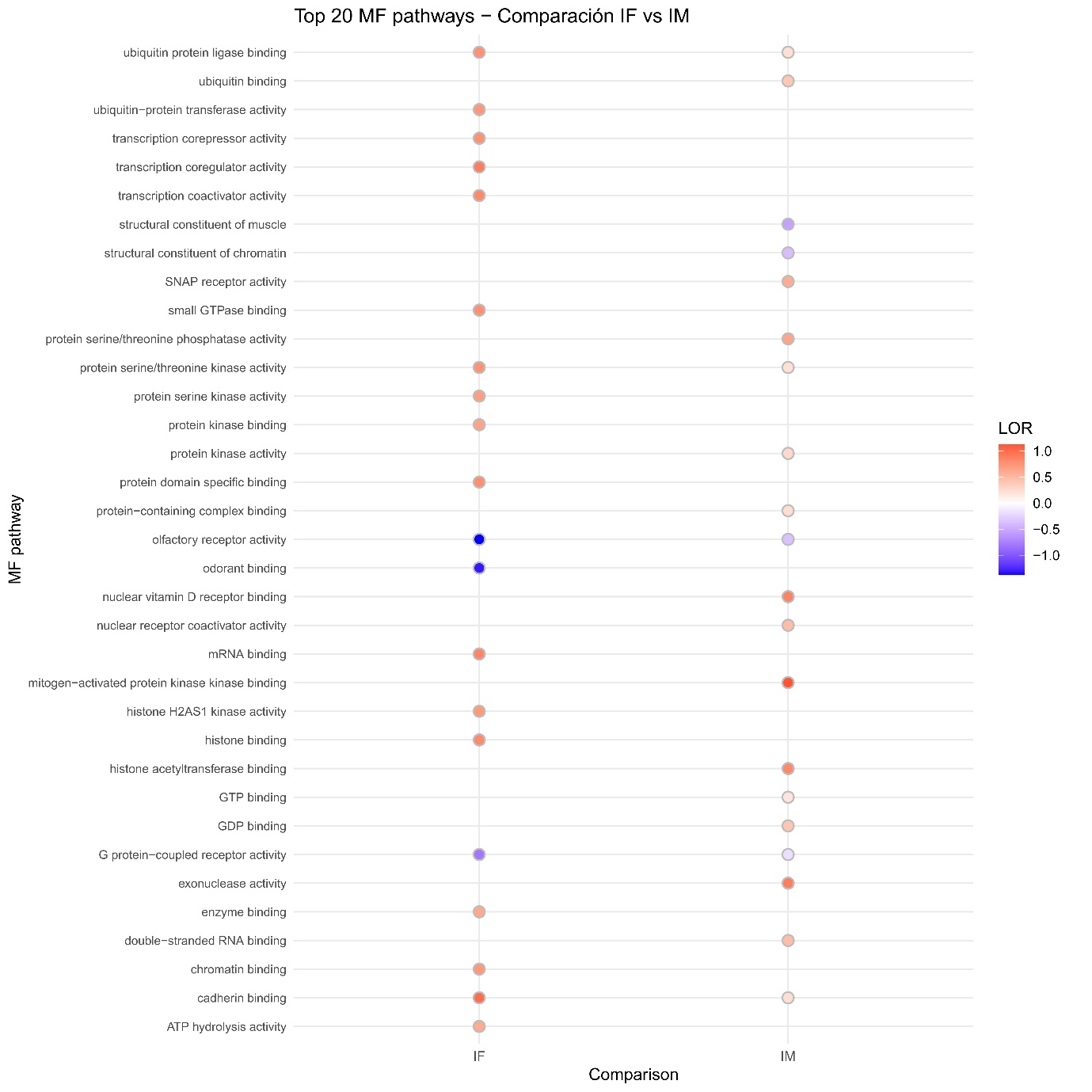
